# Differential reduction of neuropathic pain symptoms by mGlu_4_-mediated neuromodulation of amygdala circuits

**DOI:** 10.1101/2022.10.19.512850

**Authors:** Vanessa Pereira, Juri Aparicio Arias, Amadeu Llebaria, Cyril Goudet

## Abstract

Neuropathic pain is a common health problem, resulting in exacerbated response to noxious and non noxious stimuli, as well as impaired emotional and cognitive responses. Unfortunately, neuropathic pain is also one of the most difficult pain syndromes to manage, highlighting the importance of better understanding of the brain regions and neuromodulatory mechanisms involved in its regulation. Among the many interconnected brain areas which process pain, the amygdala is known to play an important role in the integration of sensory and emotional pain signals. Here, we questioned the ability of a recently identified neuromodulatory mechanisms associated to the metabotropic glutamate receptors mGlu_4_ in the amygdala to modulate neuropathic pain. In a murine model of peripheral mononeuropathy induced by a chronic constriction of the sciatic nerve, we demonstrate that pharmacological activation of amygdala mGlu_4_ receptors efficiently alleviates sensory and depressive-like symptoms in both male and female mice. Moreover, we reveal a differential modulation of those symptoms, activating mGlu_4_ receptors in the controlateral amygdala, relatively to the side of the mononeuropathy, is necessary and sufficient to relieve both sensory and depressive-like symptoms while ipsilateral activation solely reduces depressive-like symptoms. Furthermore, using photopharmacology, a recent strategy allowing a precise spatiotemporal photocontrol of deep brain endogenous targets, we further demonstrate the rapid and reversible action of mGlu_4_-mediated neuromodulation on neuropathic pain symptoms. Finally, coupling photopharmacology and analgesic conditioned place preference, we show an important pain-reducing effect of mGlu4 activation. Taken together, these data highlight the analgesic potential of enhancing amygdala mGlu_4_ activity to counteract neuropathy in the hope of improving existing treatments.

## INTRODUCTION

Neuropathic pain is caused by a lesion or disease of the somatosensory nervous system. It is a common and complex health problem, which greatly impairs the quality of life of 7-10% of the population worlwide (1, 2). Unfortunately, neuropathic pain is one of the most difficult pain syndromes to manage (3), underlining the importance of understanding the brain circuits and neuromodulatory mechanisms of pain with the hope to improve treatments.

The processing of pain-related information behind the sensory, cognitive and emotional– affective aspects of pain is very complex and involves many interconnected brain areas working together (4). Among them, the amygdala is known to be a critical region that integrates sensory and pain signals (5). The amygdala receives pain-related information mainly from the thalamus, cortical areas and the parabrachial nucleus (5–7). The amygdala is composed of multiple interconnected nuclei, comprising the basolateral (BLA) and central (CeA) nuclei and the intercalated cells (ITC), which have all been linked to pain-related functions. Within amygdala, various neuromodulatory systems are implicated in the modulation of those functions, such as opioids (8), cannabinoids (9, 10), neuropeptides (8, 11), as well as glutamate (12–14).

Glutamate is one of the main neurotransmitters involved in the transmission of pain-related information throughout the nervous system. It exerts its action via the activation of ionotropic and metabotropic receptors. Metabotropic glutamate receptors (mGlu) are G protein-coupled receptors activated by the neurotransmitter glutamate (15) and several studies have highlighted the analgesic potential of these receptors (14, 16). Recently, another member of the mGlu receptors family, the mGlu_4_ subtype, has been identified in the amygdala where it acts as a neuromodulator of sensory and anxiodepressive symptoms associated to persistent inflammatory pain (17). These receptors are present mainly in presynaptic elements of both glutamatergic and GABAergic neurons within the LA and BLA, and they downregulate the transmission coming from the thalamus (17). mGlu_4_ receptors could make interesting analgesic targets against pathological pain because, while leaving acute pain in naïve animals unchanged, systemic or local administration of mGlu_4_ agonists in the spinal cord or the amygdala alleviates pain in animal models of chronic pain (17–21). However, the role of these receptors in chronic pain from various etiologies remains to be further explored.

In the present study, we questioned the ability of amygdala mGlu_4_ receptors to modulate neuropathic pain. To that aim, we combined classical behavioral pharmacology in a mouse preclinical model of peripheral mononeuropathy and photopharmacology, a recent strategy allowing a precise spatiotemporal control of deep brain endogenous targets by light-operated ligands (22, 23).

## MATERIALS AND METHODS

### Animals

Experiments were performed on 8- to 12-week-old C57BL/6J male and female mice (Charles River). Animals were treated in accordance with the European Community Council Directive 2010/63/EU. Experimental protocols were approved by the regional animal welfare committee (CEEA-LR) with the guidelines of the French Agriculture and Forestry Ministry (D34-172-13). All efforts were made to minimize animal suffering and to reduce their number according to the 3R principles.

### Stereotaxic implantation of cannulas

Prior to neuropathy induction and behavioral testing, guide cannulas for pharmacology (PlasticsOne, Roanoke, VA) or hybrid opto/fluidic cannulas for photopharmacology (DORIC lenses, Quebec, Canada) were implanted unilaterally by stereotaxic surgery on anesthetized mice. Cannulas were placed over the intermediate capsule of right or left amygdala (−1.34 mm anteroposterior (AP); ±2.9 mm mediolateral (ML); and −4.25 mm dorsoventral (DV)). After 1 week of recovery from the stereotaxic surgery, animals were first subjected to different behavioral tests to measure their basal locomotor activity, mechanical or thermal sensitivity and grooming behavior in order to establish a baseline. At the end of the series of behavioral experiments, brains were post-fixed to check the cannula locations.

### Peripheral mononeuropathy induction

Induction was performed after 1 week of recovery from stereotaxic implantation and baseline measurements. We used the “Cuff model” of neuropathic pain, known to induce long-lasting sensory and anxiodepressive symptoms in mice (24). Peripheral mononeuropathy was induced by the unilateral implantation of a cuff made of a short polyethylene tube (2 mm) around the main branch of the sciatic nerve of the right or left hind paw of anesthetized mice (25). Sham surgeries followed the same procedure, but without implantation.

### Behavioral experiments

Mechanical sensitivity was evaluated using the von Frey method. The mechanical force required to elicit a paw withdrawal response in 50% of animals (in grams) was determined using the simplified “up-down” Von Frey method (26). Heat sensitity was measured using the Hargreaves test. A radiant infrared heat stimulus was focused on the plantar surface of the hindpaws of mice to determine the time taken (in seconds) to withdraw from the heat stimulus (27). Depressive-like behavior was assessed using the splash test (28). The duration that mice spent pursuing grooming behavior after spraying a 10% sucrose solution on their dorsal coat was recorded manually over a total period of 5 min for classical pharmacology and 9 min for photopharmacology experiments.

These behaviors were tested on healthy mice (baseline, before surgery) and on neuropathic or sham operated mice (>14 days after the surgery, as indicated in the corresponding figures) after intra-amygdala injection of drugs or their vehicle. Locomotor activity and additional behaviors were also recorded **(Supplemental Figure 1)**.

### In vivo photopharmacology

*In vivo* photopharmacology was applied in experiments measuring mechanical and heat allodynia, depressive-like behavior and in analgesic conditioned place preference experiments in mononeuropathic mice. Experiments were performed on mice stereotaxically implanted with hybrid opto/fluidic cannulas. Mice were connected *via* a catheter to a minipump and *via* an optical fiber to a LED source, allowing the local delivery of drugs and light in the amygdala in freely moving animals. We used a LED light source (DORIC lenses, Quebec, Canada) combining two wavelengths (UV: 385nm; Green: 505 nm) which are independently controlled via the LED driver software (Doric Lenses, Quebec, Canada) and connected through a rotary joint to an optical fiber (fiber diameter: 200 μm, NA = 0.53). Mice were habituated to be connected daily during one week before the tests. Tests were performed after intra-amygdala injection of optogluram or vehicle. Mice received 50 ms light pulses at 10 Hz frequency and a light power of 8.0 mW for 385 nm wavelength and 2.0 mW for 505 nm wavelength. The duration of light exposure was adapted for each behavioral test. Typically, light application started 15 minutes following injection, when the ligand reaches its maximal effect (as determined in absence of light).

### Analgesic conditioned place preference (aCPP)

In order to evaluate the analgesic potential of mGlu_4_ photocontrol in absence of external stimuli, we used the aCPP paradigm (29), combined with photopharmacology. The aCPP apparatus consists in a two-chamber arena presenting different contexts (striped or dotted wall patterns), which are connected through a central open door. One chamber was defined as the “violet chamber” and the other one as the “green chamber”. The illumination is automatically controlled through a video tracking device coupled to the light source controller (EthoVision, Noldus, Wageningen, Netherlands). When the mouse is detected in the “violet chamber”, it receives a 385 nm LED illumination in the amygdala through an optic fiber. On the other hand, when the mouse is in the “green chamber”, it receives a 505 nm LED illumination in the amygdala. Following a first session of habituation to the arena in absence of treatment, mice were submitted to 10 conditioning episodes of 5 minutes, twice daily for 5 days. During each conditioning episode, neuropathic or sham operated mice were injected with either vehicle (PBS) or optogluram (30 μM, 500nL in PBS). Fifteen minutes after injection (when drug reached its maximal effect), mice were placed for 5 minutes in the arena and allowed to move freely. Mice were first placed alternatively in one or the other chamber. The 6^th^ day, the animals were placed in the center of the arena, receiving no drug or light treatment, and their real-time place preference was measured during 5 minutes through a video tracking software.

### Ligands and chemicals

All chemicals were reagent grade (Merck, or Sigma, Germany). Optogluram and LSP4-2022 were synthesized following the experimental procedures previously reported (30, 31).

### Statistics

All data are reported as mean ± standard error of the mean (SEM). Number of mice and statistical tests that were performed on datasets are indicated in Figure Legends. Data were analyzed using Prism software (GraphPad, La Jolla, CA, USA) using one-way or two-way analysis of variance (ANOVA) and the appropriate post-hoc tests for multiple comparisons. Data were considered significant when p<0.05.

## RESULTS

### Peripheral mononeuropathy induces mechanical and heat allodynia and depressive-like symptoms in male and female mice

Along this study, we used a preclinical model of peripheral mononeuropathy, named the “cuff model”, provoked by the unilateral implantation of a tube enclosing the sciatic nerve. In mice, this model is known to induce long-lasting sensory symptoms, such as mechanical and heat allodynia, as well as anxiodepressive-like symptoms, such as a defect in grooming behavior (24). This model reproduces similar symptoms to those of patients with neuropathic pain who often suffer from allodynia, a pathological state in which an innocuous stimuli, becomes painful (3) as well as depression which is one of the most common comorbidities of neuropathic pain (1).

We measured mechanical and thermal sensitivity, as well as grooming behavior before surgery (Baseline, D0) and two to three weeks after the induction **(Figure 1a)**. After 14 days, we observed a significant decrease in the paw withdrawal threshold on the side of the lesion, as measured by the Von Frey technique (**Figure 1b, e)**. We also observed a significant decrease of the latency to withdraw the paw from heat source, as measured by the Hargreaves test **(Figure 1c, f)**. This indicates that the chronic constriction of the sciatic nerve provoked by the cuff induces both mechanical and heat allodynia. In addition, mice elicit a significantly reduced duration of grooming behavior when submitted to the splash test, indicative of depressive-like behavior **(Figure 1c)**.

**Figure 1.**
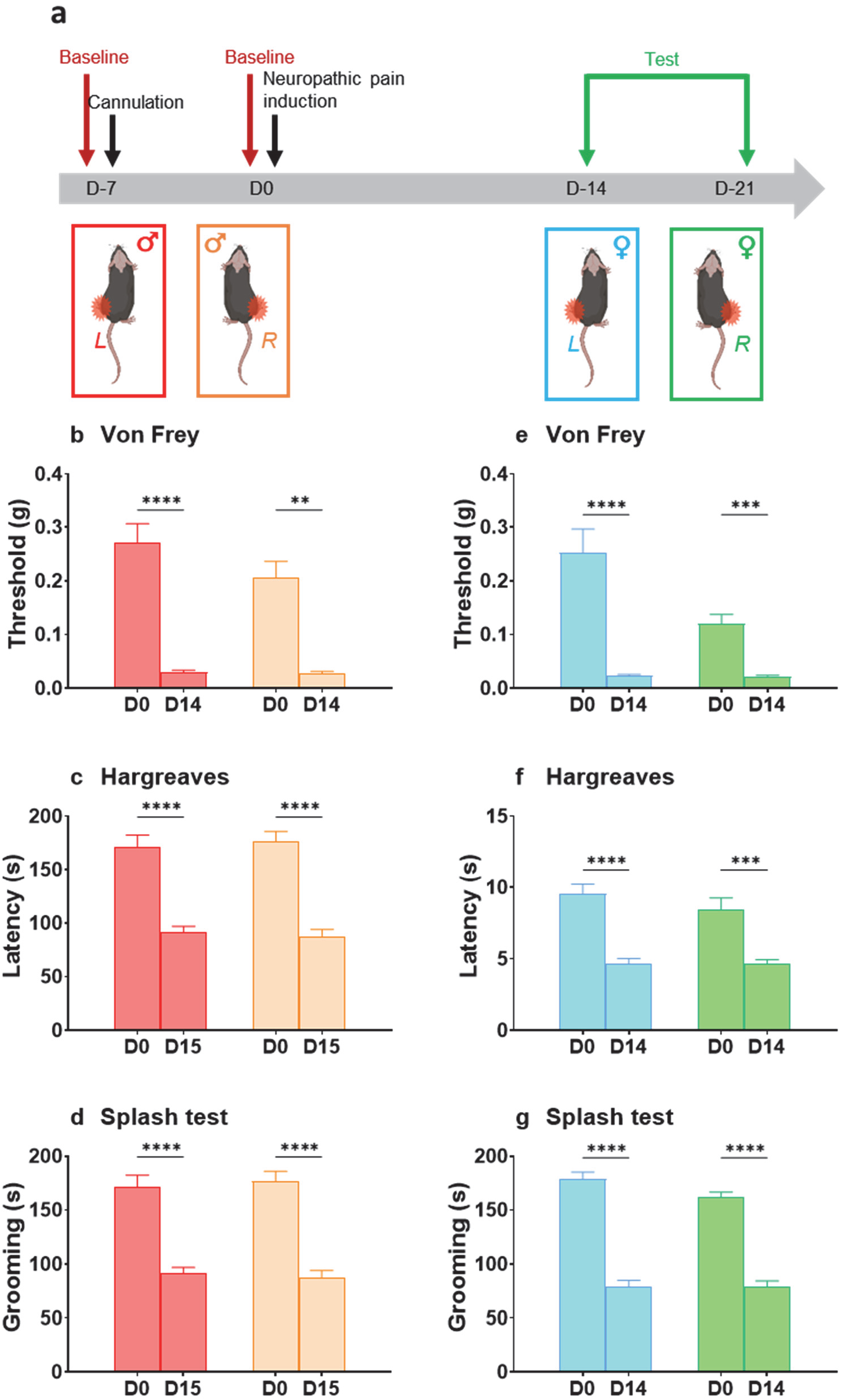
Cuff-induced mononeuropathic pain model leads to sensory and depressive-like symptoms in male and female mice symptoms. Peripheral mononeuropathy was induced by positioning a cuff around the sciatic nerve of the left or right hind paw of male and female mice. **a**: Experimental design and timeline of behavioral tests. **b, e:** Mechanical allodynia was observed both in male or female mice, 14 days after implantation of the Cuff either on the left or right hind paw, as indicated by the significant reduction of the ipsilateral paw withdrawal thresholds measured using the Von Frey technique (n=10, D0 vs D14, two-way ANOVA, Sidak’s post-hoc test). **c, f:** Heat allodynia was observed both in male or female mice, 14 days after implantation of the Cuff either on the left or right hind paw, as indicated by the significant reduction of the latency to withdraw the paw in the Hargreaves test (n=10, D0 vs D14, two-way ANOVA, Sidak’s post-hoc test). **d, g:** Depressive-like behavior assessed in the splash test revealed a significant decrease in grooming duration in both male and female Cuff mice, 15 days after induction (male n=10, female n=10, D0 vs D14, two-way ANOVA, Sidak’s post-hoc test).

### Activation of mGlu_4_ in the amygdala relieves neuropathic pain symptoms in male mice

We then questioned the ability of amygdala mGlu_4_ to modulate symptoms of neuropathic pain. To that aim, we first used a selective mGlu_4_ agonist, LSP4-2022 (31), that we injected unilaterally (5 μM, 500 nL in PBS) either in the ipsilateral or controlateral amygdala as compared to the peripheral mononeuropathy on the right or left hind paw.

We first assessed the potential modulation of neuropathic pain symptoms by right or left amygdala mGlu_4_ when the peripheral mononeuropathy is on the left hind paw. Interestingly, we observed that activation of the right amygdala mGlu_4_ significantly reduces mechanical or heat allodynia whereas activation of left amygdala mGlu_4_ does not modify them (**Figures 2a and 2b**). On the other hand, activation of mGlu_4_ in both right or left amygdala significantly restores grooming behaviour in those mice (**Figure 2c**).

**Figure 2:**
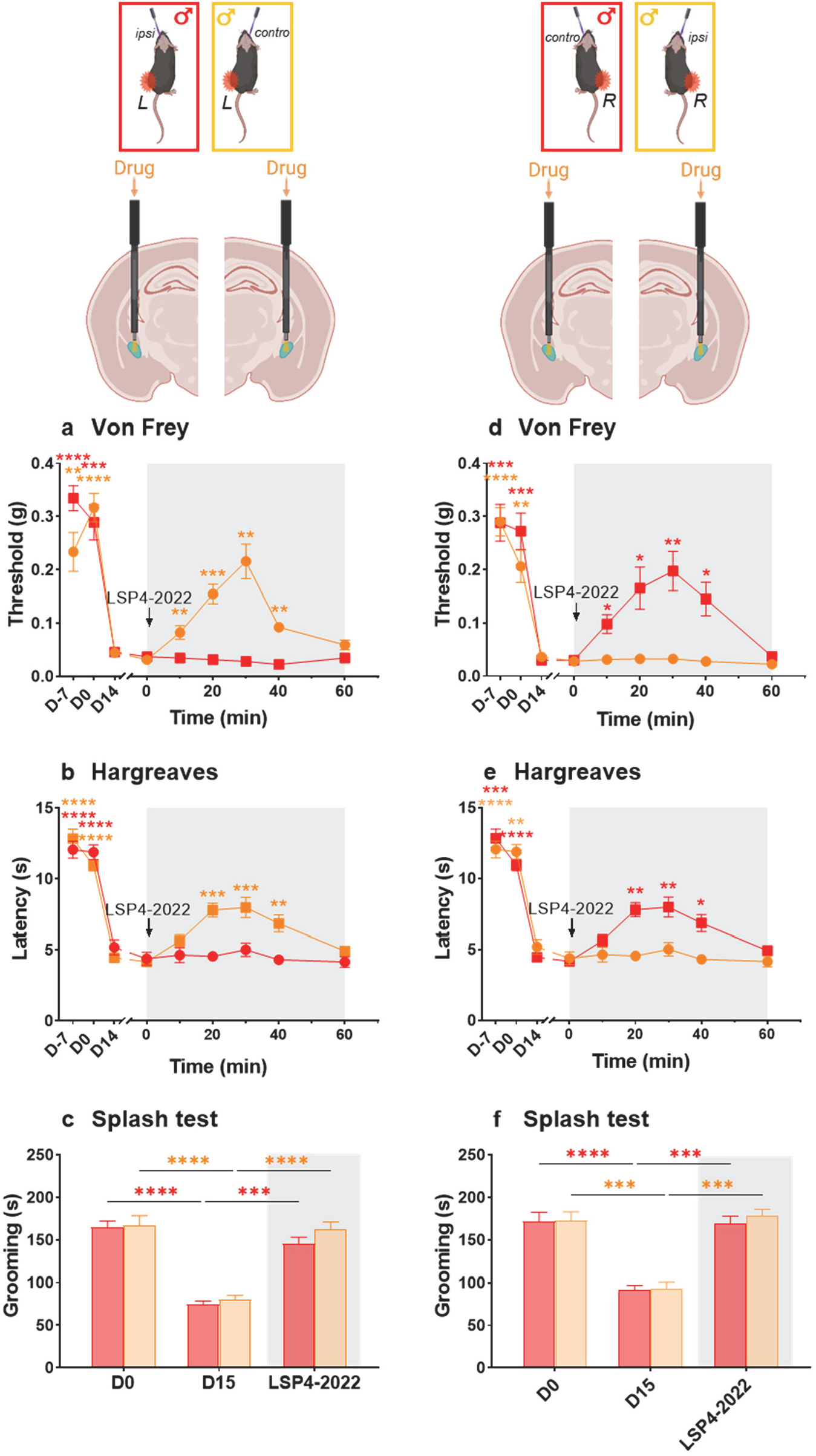
mGlu_4_ activation in the amygdala differentially inhibits hypersensitivity and depressive-like behaviour in male neuropathic mice. Peripheral mononeuropathy was induced by positioning a cuff around the sciatic nerve of the left or right hind paw of male mice. Beforehand, mice were implanted with a cannula on the ipsilateral or contralateral amygdala (left cuff for females). On the test day, at t_0_, mice were injected with the selective mGlu_4_ agonist, LSP4-2022 (5 μM, 500 nL in PBS) either on the ipsi or controlateral amygdala to the mononeuropathy and the subsequent effects on neuropathic pain symptoms were measured. **a, d:** Effect on mechanical allodynia of mGlu_4_ activation on the ipsilateral or contralateral amygdala on male mice with a mononeuropathy on the left or right hind paw. Mechanical allodynia as determined by the Von Frey technique. Mean threshold ± SEM (g), t_x_ vs t_0_ (0 min), two-way ANOVA, Dunnett’s post-hoc test..**b, e:** Effect on heat allodynia of mGlu_4_ activation on the ipsilateral or contralateral amygdala on male mice with a mononeuropathy on the right or left hind paw. Thermal allodynia as determined by Hargreaves method. Mean threshold ± SEM (s), t_x_ vs t_0_ (0 min), two-way ANOVA, Dunnett’s post-hoc test **c, f:** Effect on depressive-like symptoms of mGlu_4_ activation on the ipsilateral or contralateral amygdala on male mice with a mononeuropathy on the right or left hind paw. Depressive-like symptoms as determined by the Splash Test. Mean grooming time ± SEM; t_x_ vs t_0_ (0 min), two-way ANOVA, Tukey’s post-hoc test.

To verify whether this asymmetrical modulation of allodynia could result from a specialization of the right amygdala in the modulation of certain pain-related functions as previously observed (32), we performed a series of mirror experiments in which the peripheral mononeuropathy is on the right hind paw this time. When the neuropathy is on the right side, activation of the left amygdala mGlu_4_ significantly reduces mechanical or heat allodynia whereas activation of right amygdala mGlu_4_ does not modify them (**Figures 2d and 2e**). Activation of mGlu_4_ in both right or left amygdala significantly restores grooming behaviour in those mice (**Figure 2f**).

This indicates that activation of mGlu_4_ in the amygdala controlateral to the lesion is necessary and sufficient to relieve both sensory and depressive symptoms of peripheral mononeuropathy in male mice, while activation of mGlu_4_ in the ipsilateral amygdala solely abolishes depressive-like behavior.

Another interesting observation is that activation of amygdala mGlu_4_ by local injection of LSP4-2022 does not significantly modify mechanical or heat sensitivity, as well as grooming behavior, in Sham operated mice **(supplemental figure 2)**. As previously reported (17, 19–21), it seems that mGlu_4_ activation solely restores hypersensitivity or abnormal behaviors associated to pathological states without significantly affecting normal sensitivity or behavior in healthy mice.

### Activation of mGlu_4_ in the amygdala also relieves neuropathic pain symptoms in female mice

Chronic pain exhibits a higher prevalence in women than in men (1). Over the past years, preclinical studies have highlighted several sexual dismorphism in pain mechanisms (33–35), notably at the amygdala level (36).,Thus, we sought to determine whether amygdala mGlu_4_ activation can also modulate neuropathic pain symptoms in female mice. To that aim, we assessed the potential modulation of neuropathic pain symptoms following activation of mGlu_4_ by the agonist LSP4-2022 in the right or left amygdala on female mice suffering from peripheral mononeuropathy in their left hind paw.

First, as observed in male mice, the chronic constriction of the sciatic nerve provokes mechanical and thermal allodynia, as well as depressive-like symptoms in female mice. Then, following local injection of LSP4-2022, we obtained similar results than in males: i) controlateral activation is required to significantly diminish mechanical or heat allodynia while ipsilateral activation does not modify these sensory symptoms (**Figures 3a and 3b**) and ii) either ipsi or controlateral activation significantly restore grooming behaviour in female mice (**Figure 3c**).

**Figure 3:**
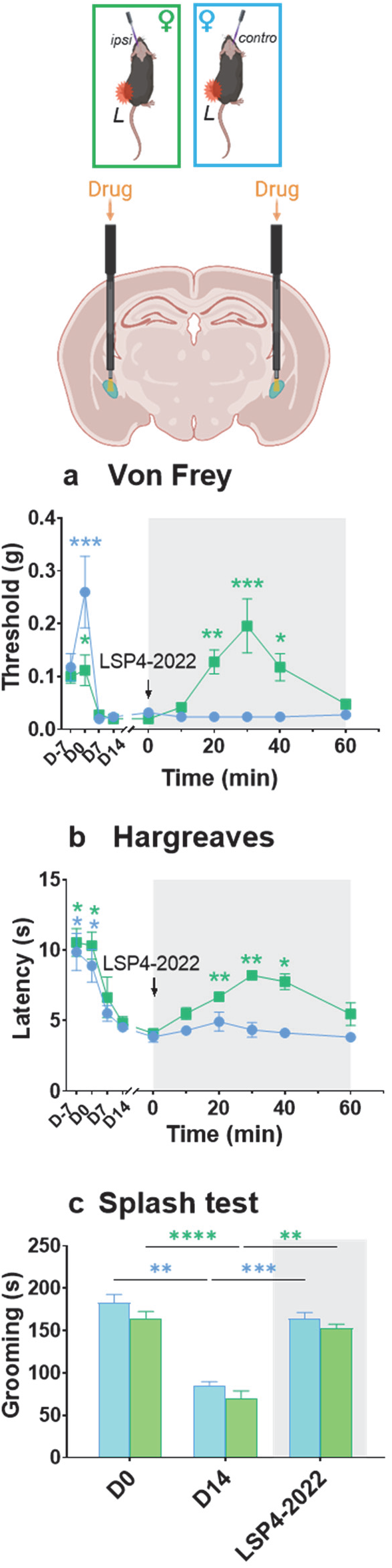
mGlu_4_ activation in the amygdala differentially inhibits hypersensitivity and depressive-like behaviour in female neuropathic mice. Peripheral mononeuropathy was induced by positioning a cuff around the sciatic nerve of the left hind paw of female mice. Beforehand, mice were implanted with a cannula on the ipsilateral or contralateral amygdala. On the test day, at t_0_, mice were injected with the selective mGlu_4_ agonist, LSP4-2022 (5 μM, 500 nL in PBS) either on the ipsi or controlateral amygdala to the mononeuropathy and the subsequent effects on neuropathic pain symptoms were measured. **a:** Effect on mechanical allodynia of mGlu_4_ activation on the ipsilateral or contralateral amygdala on female mice with a mononeuropathy on the left or right hind paw. Mechanical allodynia as determined by the Von Frey technique. Mean threshold ± SEM (g), t_x_ vs t_0_ (0 min), two-way ANOVA, Dunnett’s post-hoc test..**b:** Effect on heat allodynia of mGlu_4_ activation on the ipsilateral or contralateral amygdala on female mice with a mononeuropathy on the right or left hind paw. Thermal allodynia as determined by Hargreaves method. Mean threshold ± SEM (s), t_x_ vs t_0_ (0 min), two-way ANOVA, Dunnett’s post-hoc test **c:** Effect on depressive-like symptoms of mGlu_4_ activation on the ipsilateral or contralateral amygdala on female mice with a mononeuropathy on the right or left hind paw. Depressive-like symptoms as determined by the Splash Test. Mean grooming time ± SEM; t_x_ vs t_0_ (0 min), two-way ANOVA, Tukey’s post-hoc test.

This series of experiments demonstrates that activation of amygdala mGlu_4_ relieves neuropathic pain symptoms in both male and female mice.

### Photocontrolled-activation of amygdala mGlu_4_ dynamically alleviates neuropathic pain symptoms

Next, we used photopharmacology, an emerging strategy to control the biological activity of endogenous proteins by light, to assess whether amygdala mGlu_4_ can exert a dynamic modulation of neuropathic pain-related symptoms. This technique allows a precise spatiotemporal control of deep brain endogenous targets by light-operated ligands (22, 23).

To photocontrol amygdala mGlu_4_ receptors, we used optogluram, a photoswitchable positive allosteric modulator (PAM) of mGlu_4_ receptors (17, 30). Optogluram posseses an azobenzene moiety in its scaffold, acting as a chemical switch. Optogluram is active in the dark then, under illumination with violet light, it isomerizes from its active trans-isomer to its inactive cis-isomer. Under illumination with green light, it switches back to its trans active isomer (17). Experiments were performed on mice implanted with hybrid optic-fluidic cannula connected to a minipump for drug injection and to a LED source through optic fibers for illumination. After a basal measurement at t_0_, ligand or vehicle were injected unilaterally either in the ipsilateral or controlateral amygdala as compared to the peripheral mononeuropathy on the left hind paw. After 15 minutes, the mechanical and heat sensitivity is measured to determine the effect of the treatment on these parameters. Then, from 15 to 45 minutes, we applied 3 successive cycles of violet/green illumination to deactivate/reactivate optogluram and measured mechanical or heat sensitivity after each illumination period of 5 minutes.

In the dark, neuropathic-induced mechanical or heat allodynia are abolished 15 minutes after contralateral intra-amygdala microinjection of optogluram (30μM in 500nL of PBS), but not following ipsilateral injection. As can be seen in **Figure 4a and 4b**,.the antiallodynic action of optogluram is switched off following a 5 minutes illumination with violet light (λ=385nm, 8.0mW, 10Hz) and recovered after 5 minutes of green illumination (λ=505nm, 2.0mW, 10Hz). Intra-amygdala injection of optogluram also reduced the depressive-like behavior of neuropathic mice measured with the splash test. In the dark, 15 minutes after ipsi or contralateral administration, optogluram (30μM in 500nL of PBS) increased the grooming duration of cuff-mice, whereas 3 minutes of violet light illumination (λ=385nm, 8.0mW, 10Hz) reduced it to the levels observed in vehicle-treated cuff mice. Then, the increase of the grooming duration is restored following 3 minutes of green light illumination (λ=505nm, 2.0mW, 10Hz) **(Figure 4c)**. We verified that light by itself doesn’t modify the different symptoms measured in animals injected with saline solution (500 nL of PBS). Violet- or green-light illumination in the amygdala had no effect on the measured parameters in absence of the photoswitchable mGlu_4_ PAM **(supplemental figure 3**). Also, photopharmacological manipulation of amygdala mGlu_4_ using optogluram does not modify mechanical sensitivity, thermal sensitivity or grooming in Sham operated mice **(supplementary figure 4)**, similar to what was observed following mGlu_4_ activation with the agonist LSP4-2022 **(supplemental figure 2)**.

**Figure 4:**
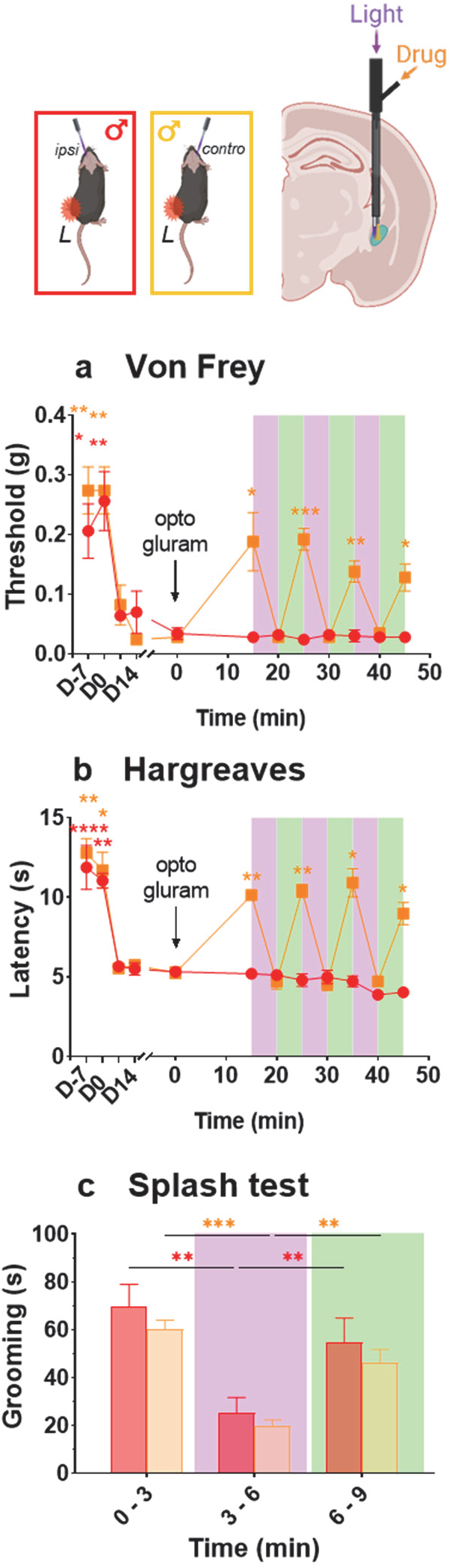
Photopharmacological manipulation of mGlu_4_ in the amygdala differentially inhibits hypersensitivity and depressive-like behaviour of neuropathic mice. Effects of amygdala mGlu_4_ photocontrol on neuropathic pain symptoms were measured on male mice with a cuff implanted around the left hind paw, from 14 to 21 days post induction Local drug and light delivery was performed through a stereotaxically implanted hybrid optofluidic cannula. On the test day, at t_0_, mice were injected with the photoswitchable mGlu_4_ enhancer, optogluram (30 μM, 500 nL in PBS) either on the ipsi or controlateral amygdala to the mononeuropathy. Violet (385 nm, 10 Hz, 8 mW) or green light (505 nm, 10Hz, 2 mW) was applied by the mean of an optic fiber connected to a LED light source and a controller. **a**. Mechanical allodynia as determined by the Von Frey technique. Mean threshold ± SEM (g), t_x_ vs t_0_ (0 min), two-way ANOVA, Dunnett’s post-hoc test. **b**. Thermal allodynia as determined by Hargreaves method. Mean threshold ± SEM (s), t_x_ vs t_0_ (0 min), two-way ANOVA, Dunnett’s post-hoc test. **c**. Depressive-like symptoms as determined by the Splash Test. Mean grooming time ± SEM; t_x_ vs t_0_ (0 min), two-way ANOVA, Tukey’s post-hoc test.

Taken together, these results demonstrate the dynamic nature of mGlu_4_ control over neuropathic pain-related symptoms.

### Amygdala mGlu_4_ photocontrolled activation promotes analgesic condititioned place preference

Next, we used the analgesic conditioned place preference paradigm (aCPP)(29) to evaluate the analgesic potential of amygdala mGlu_4_ activation in neuropathic mice, without the involvement of external noxious stimuli. For an enhanced precision, we coupled aCPP with photopharmacology.

Experiments were performed in right implanted male mice, while the mononeuropathy was on the left hind paw. Experimental conditions for photopharmacological manipulations were similar to those described above, except that illumination was automatically controlled through a videotracking device detecting the location of the mouse.

Conditioning was performed twice daily for 5 days. During this conditioning period, neuropathic or sham mice received an intra-amygdala administration of either vehicle (500 nL of PBS) or optogluram (30μM in 500nL of PBS) and, depending on their position in a two-chamber arena, they automatically received either a green or violet illumination through an optic fiber **(Figure 5a, b)**. We verified that chronic treatment with optogluram does not lead to tolerance. Neuropathic mice received an intra-amygdala microinjection of optogluram (30μM in 500nL PBS) twice daily for 5 days, which corresponds to the conditioning period. We measured the antiallodynic effect of optogluram on day 6 **(supplementary figure 5)** and observed no difference on the peak effect of the drug (15 minutes following intra-amygdala injection of optogluram) on the paw withdrawal threshold measured by the Von Frey technique that was measured at D14 post-induction.

**Figure 5:**
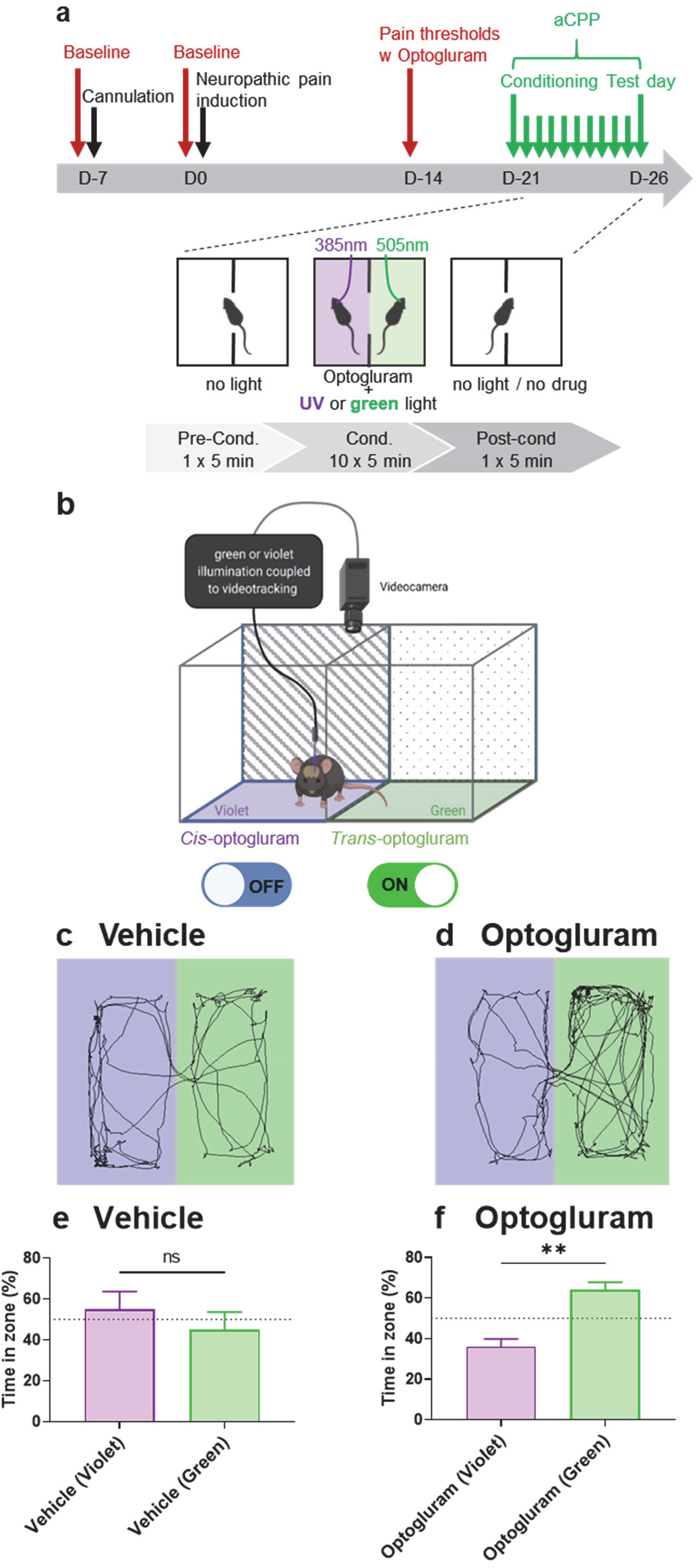
Photopharmacological manipulation of amygdala mGlu_4_ promotes analgesic conditioned place preference (aCPP) of neuropathic mice. **a:** Experimental design and timeline of behavioral tests. On these experiments, neuropathy was induced by application of a cuff on the left hind paw of male mice, beforehand implanted with a hybrid optofluidic cannula on the contralateral (right) amygdala. The conditioning took place between from day 21 to day 25, the test day was day 26 post surgery. Mice were submitted to 10 conditioning episodes of 5 minutes, twice daily for 5 days. During each conditioning episode, animals were injected with either vehicle (PBS) or optogluram (xxx μM, 500nL in PBS). Fifteen minutes after injection (when drug reached its maximal effect), mice were placed for 5 minutes in a two-chambers arena and allowed to move freely. Mice were first placed alternatively in the one or the other chamber. The 6^th^ day, the animals were placed in the center of the arena, receiving no drug or light treatment, and their real-time place preference was measured during 5 minutes through a videotracking software. **b:** Schematic representation of the aCPP setup. The arena consists in two-chambers with different context (striped or dotted walls) connected through a central open door. The illumination is automatically controlled through a video tracking device coupled to the light source controller. When the mouse is detected in the “violet chamber”, it receives a 385 nm LED illumination, while it receives a 505 nm LED illumination when it is in the green chamber. **c, d:** Test results: representative 5-minutes tracks of a neuropathic mouse which received either vehicle (c) or optogluram (d) during the conditioning period. **e, f:** Test results: mean percentage ± SEM of time spent in the “green chamber” or the “violet chamber” of neuropathic mice treated with Vehicle (n=8) or with Optogluram (n=12) during conditioning (Mean ± SEM, Time in Violet vs Time in Green area, one-way ANOVA, Tukey’s post-hoc test).

On the 6^th^ day, we tested the eventual preference of neuropathic or sham mice for the “green chamber” (in which optogluram was switched-on through green illumination) over the “violet chamber” (in which it was switched-off through violet illumination). During this test, mice received no drug or light treatment. The time spent in each chambers is measured using the videotracking device. As can be seen in **Figure 5c-f**, the group of conditioned neuropathic mice that received the optogluram treatment significantly preferred the green area, contrary to the conditioned mice that received saline which exhibited no preference. No difference between the time spent in the violet or green areas was observed in non-conditioned Sham or neuropathic mice **(supplementary figure 6 and 7).**

These experiments demonstrate the analgesic potential of amygdala mGlu_4_ activation in neuropathic mice.

## DISCUSSION

In this study, we demonstrate that sensory and depressive symptoms of neuropathic pain are rapidly relieved under mGlu_4_ control in male and female mice, and that ipsi or controlateral amygdala differentially contribute to the modulation of these symptoms. The controlateral amygdala is necessary and sufficient to alleviate both sensory and depressive-like symptoms resulting from peripheral mononeuropathy, while mGlu_4_ activation in the ipsilateral amygdala is only reducing depressive-like symptoms **(Figure 6)**. Using photopharmacology, we reveal that amygdala mGlu_4_ exerts a rapid and dynamic control over neuropathic pain-related symptoms. The analgesic potential of amygdala mGlu_4_ activation was further underlined in the conditioned place preference paradigm. This method has been succesfully used to probe the efficacy of various analgesics (29, 37). The interest of aCPP is that subject animals determine by themselves the analgesic efficacy of a given treatment, without the involvement of external noxious stimuli. Here, we coupled aCPP with photopharmacology for the first time to our knowledge. All mice received a similar treatment by a photoswitchable mGlu_4_ enhancer and, depending on their position in a two-chamber arena automatically detected by a videotracking device coupled to the illumination source, the ligand was activated or deactivated by light. After the conditioning period, mice clearly preferred the context in which the activity of amygdala mGlu_4_ receptors was potentiated. Also of interest, we did not observe a decrease efficacy of antiallodynic action following repeated treatment by optogluram, indicating a lack of tolerance **(supplemental figure 5)**. Our results extend previous works demonstrating that exogenous activation of mGlu_4_ receptors is benificial for chronic pain symptoms. Systemic administration of the mGlu_4_ selective agonist LSP4-2022 relieves symptoms of chronic pain from different etiologies (20). At the spinal cord level, mGlu_4_ receptors are found in the inner laminae II of the dorsal horn, a region that receives the afferences from nociceptive Aδ- and C-fibers, where its activation reduces excitatory neurotransmission through the inhibition of Ca^2+^ entry via N or P/Q type voltage-gated calcium channels in the presynaptic terminal (20). Intrathecal injection of mGlu_4_ agonists, such as LSP4-2022, or allosteric enhancers, such as PHCCC or VU0155041, alleviates allodynia and hyperalgesia induced by both inflammatory or neuropathic pain without altering acute pain perception in naive animals (19–21). At the supraspinal level, mGlu_4_ receptors have been identified in important regions for pain processing, such as the thalamus (38) and the amygdala (17). Bilateral activation of amygdala mGlu_4_ alleviates pain symptoms in a mouse model of persistent inflammatory pain (17). Taken together, these results demonstrate the ability of mGlu_4_ to modulate various symptoms of chronic pain of different etiologies, without modifying acute pain perception, reinforcing its therapeutic interest for the treatment of pathological pain.

**Figure 6:**
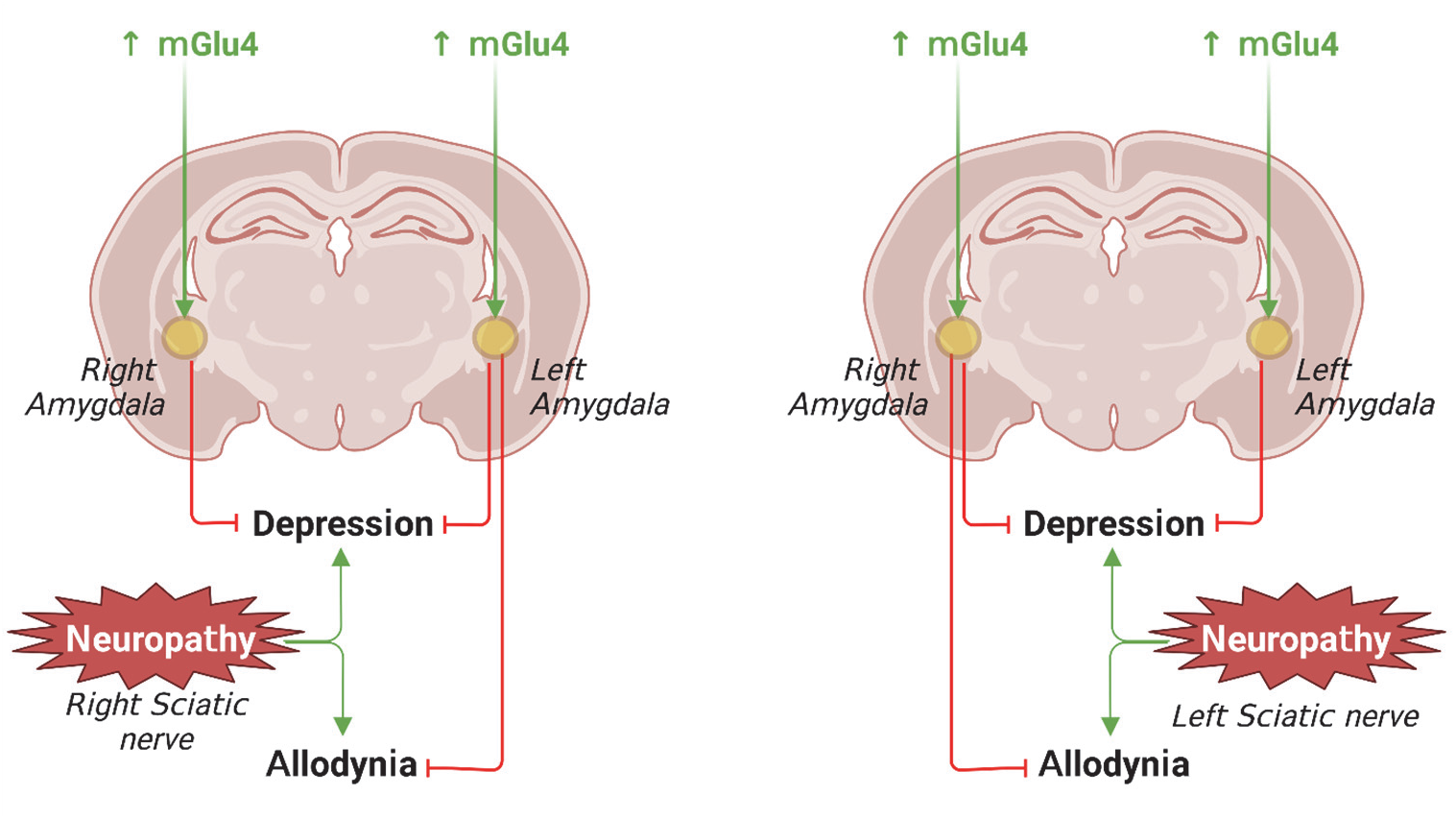
Controlateral amygdala mGlu_4_ is necessary and sufficient to relieve both sensory and depressive-like symptoms of peripheral mononeuropathy. Left hindpaw neuropathic pain sensory and depressive symptoms are relieved by right amygdala mGlu_4_ activation whereas left amygdala mGlu_4_ activation solely abolishes depressive-like symptoms. Conversely, all symptoms resulting from right hindpaw neuropathic pain are relieved by left amygdala mGlu_4_ activation whereas right amygdala mGlu_4_ activation solely abolishes depressive-like symptoms. Ipso facto, activation of mGlu_4_ in either ipsilateral or controlateral amygdala to the mononeuropathy abolishes mice depressive-like behavior.

Curiously, we did not observe an asymmetrical lateralization between the right or left amygdala. Indeed, previous studies have described a hemispheric specialization of pain-related functions in amygdala (see (32) for review). For example, independently of the inflamed side, the modulation of mechanic allodynia solely occurs following the blockade of right but not left CeA mGlu5 receptors and ERK pathways (13, 39, 40). Similarly, sensitization is only observed in the right CeA following right or left monoarthritis (41). In our experiments however, mGlu_4_ receptors from both sides can modulate sensory or depressive-like symptoms, depending on their relative position to the peripheral mononeuropathy, the key for the modulation of sensory symptoms being the localization on the contralateral side to the constriction of the sciatic nerve. Noteworthy, studies revealing the specialization of amygdala function in pain were performed in CeA (32), whereas the amygdalar expression of mGlu_4_ receptors is mainly restricted to terminals arriving in the BLA and the LA, with only little or no expression in the CeA (17). Thus, the absence of specialization of left or right amygdala mGlu_4_ function could result from the localization of mGlu_4_ receptors in LA and BLA rather than in CeA.

Interestingly, the fact that amygdala mGlu_4_ on the ipsi or controlateral side to the nerve constriction differentially contributes to pain modulation suggests that mGlu_4_ achieves its analgesic effects through the neuromodulation of different circuits **(Figure 6)**. Indeed, while mGlu_4_ activation in the controlateral amygdala to the mononeuropathy relieves both sensory and depressive-like symptoms, their activation on the ipsilateral amygdala solely decreases depressive-like symptoms. This indicates that at least two different circuits are at play, differentially regulating sensory and anxiodepressive components, and that mGlu_4_ can modulate both of them. These circuits remain to be identified.

The necessity to activate mGlu_4_ on the contralateral side to the mononeuropathy to alleviate hypersensitivity suggests that the regulation of sensory symptoms may occur through a modulation of the sensory modalities coming from thalamus nuclei (5). Indeed, in the spinothalamic tract, most secondary projection neurons of the spinal cord which transmit nociceptive information received from peripheral sensory neurons decussate and send ascending information terminating in various thalamic nuclei (4). As a result, activation of the thalamus is significantly greater in the hemisphere contralateral to the stimulus, consistent with its involvement in the processing the sensory-discriminative aspects of pain (42). This means that peripheral nociceptive information from the left side of the body is transmitted to the right side at the supraspinal level, and conversely. We have previously shown that mGlu_4_ receptors are expressed in presynaptic terminals of glutamatergic and GABAergic neurons arriving in the LA and BLA, where they downregulate the transmission coming from the thalamus (17). Their activation could in turn normalize the activities of LA and BLA neurons. One of the main target of those neurons is CeA. Thus, we can speculate that mGlu_4_ may modulate the activity within the BLA–CeA circuit, which has been implicated in the generation and modulation of pain-like behaviors (5, 43). Besides the CeA, several pathways regulating specific aspects of pain originating from BLA have been identified recenly. For example, the BLA-mPFC-PAG pathway is crucial for the development of mechanical and thermal hypersensitivity after peripheral nerve injury (44) and could be another pathway involved in the reduction of mechanical and thermal hypersensitivity following mGlu_4_ activation. We can also hypothesized that mGlu_4_ activation could modulate the neural ensemble within BLA identified by Corder and colleagues that mediates chronic pain unpleasantness (45). However, these points remain largely speculative and further studies will be required in order to identify the input and output circuits modulated by amygdala mGlu_4_ receptors.

In conclusion, this study provides strong evidence for a rapid and reversible regulation of neuropathic pain following activation of mGlu_4_ receptors in the amygdala. The data underline the therapeutic potential of mGlu_4_ receptors for chronic pain management.

## FUNDING AND DISCLOSURE (COVERS ALL AUTHORS AND SOURCES OF FUNDING LISTED FOR THE MANUSCRIPT)

This work was supported by the International Emerging Action (IEA) from the Centre National de la Recherche Scientifique (CNRS) to CG and AL and by grants from the French National Research Agency (ANR): ANR-16-CE16-0010 to CG and ANR-17-NEU3-0001 under the frame of Neuron Cofund to CG and AL.

## ACKNOWLEDGMENTS (INCLUDES SPECIAL THANKS OR DEDICATIONS)

The authors wish to thank Dr. Francine Acher for kindly providing LSP4-2022, Dr. Guillaume Lebon, Dr. Julie Le Merrer, Dr. Jérôme Becker and Dr. Laurent Givalois for helpful discussions and Dr. Ebba L. Lagerqvist for critical reading of the manuscript. Behavioral experiments were performed at the *iExplore* facility of Montpellier Biocampus and the authors thank the staff of the facility for their technical help. The illustrative graphics were created with BioRender.com.

## AUTHOR CONTRIBUTIONS (MANDATORY) FOR ALL AUTHORS

VP and CG conceived the original idea and contributed to conception of the study. VP, JAA, and CG designed the experiments. VP and JAA performed experiments. VP, JAA and CG analyzed the data. AL contributed essential materials. All authors contributed to interpretation of the results. CG wrote the manuscript and all authors provided critical feedback on the manuscript.

## SUPPLEMENTAL FIGURES AND LEGENDS

**Supplemental figure 2:**
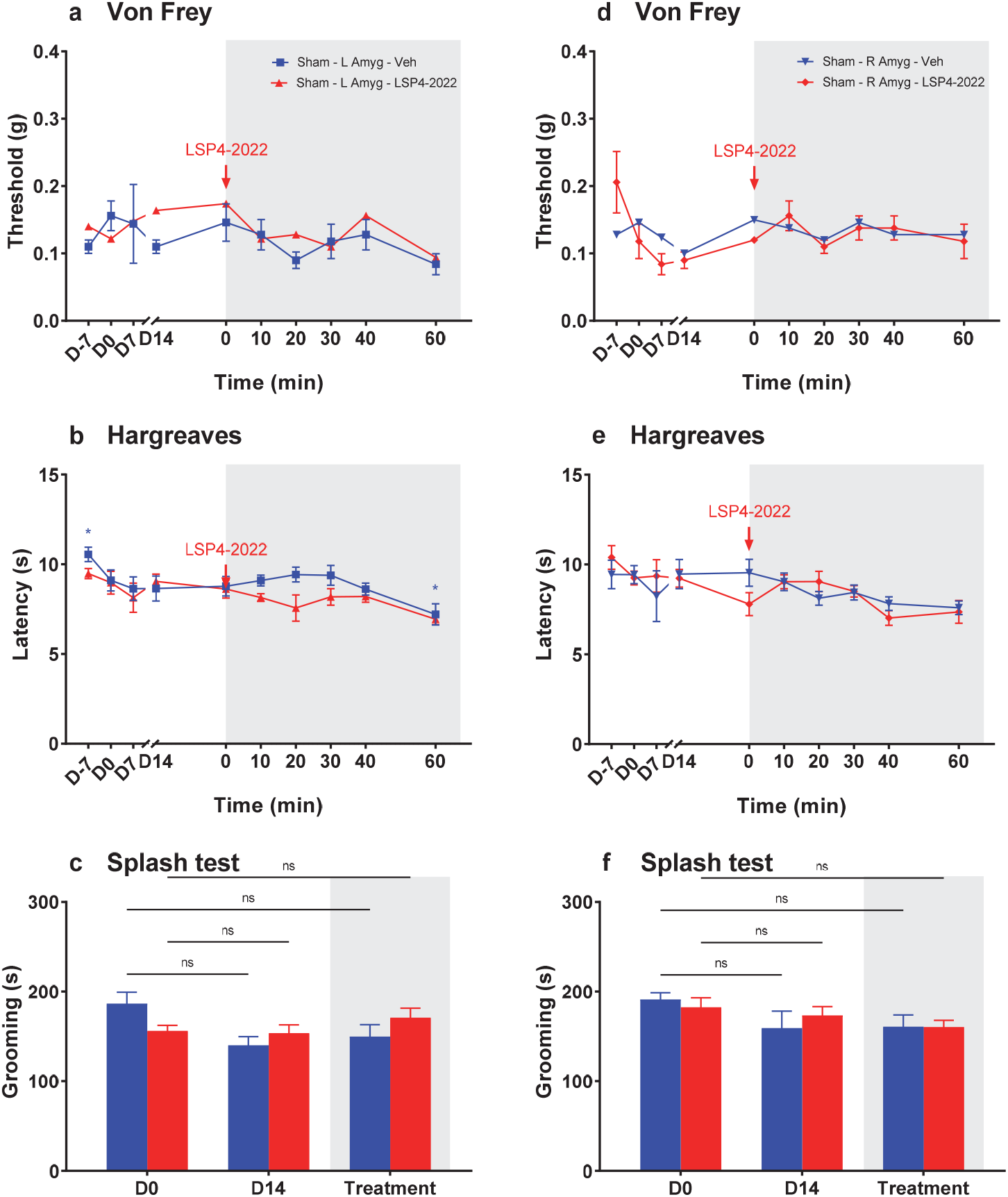
mGlu_4_ activation in right or left amygdala does not modify mechanical sensitivity, heat sensitivity or grooming in Sham animals. **a-f:** Females, sham left hindpaw, Vehicle (500 nL PBS, blue) or LSP4-2022 (5 μM, 500 nL in PBS, red) delivered unilaterally in ipsi or controlateral amygdala. **a, d**. Von Frey. Mean ± SEM, t_x_ vs t_0_ (0 min), two-way ANOVA, Dunnett’s post-hoc test. **b, e**. Hargreaves. Mean ± SEM, t_x_ vs t_0_ (0 min), two-way ANOVA, Dunnett’s post-hoc test.. **c, f**. Splash Test. # t_x_ vs t_0_ (0 min), two-way ANOVA, Tukey’s post-hoc test.

**Supplemental Figure 3:**
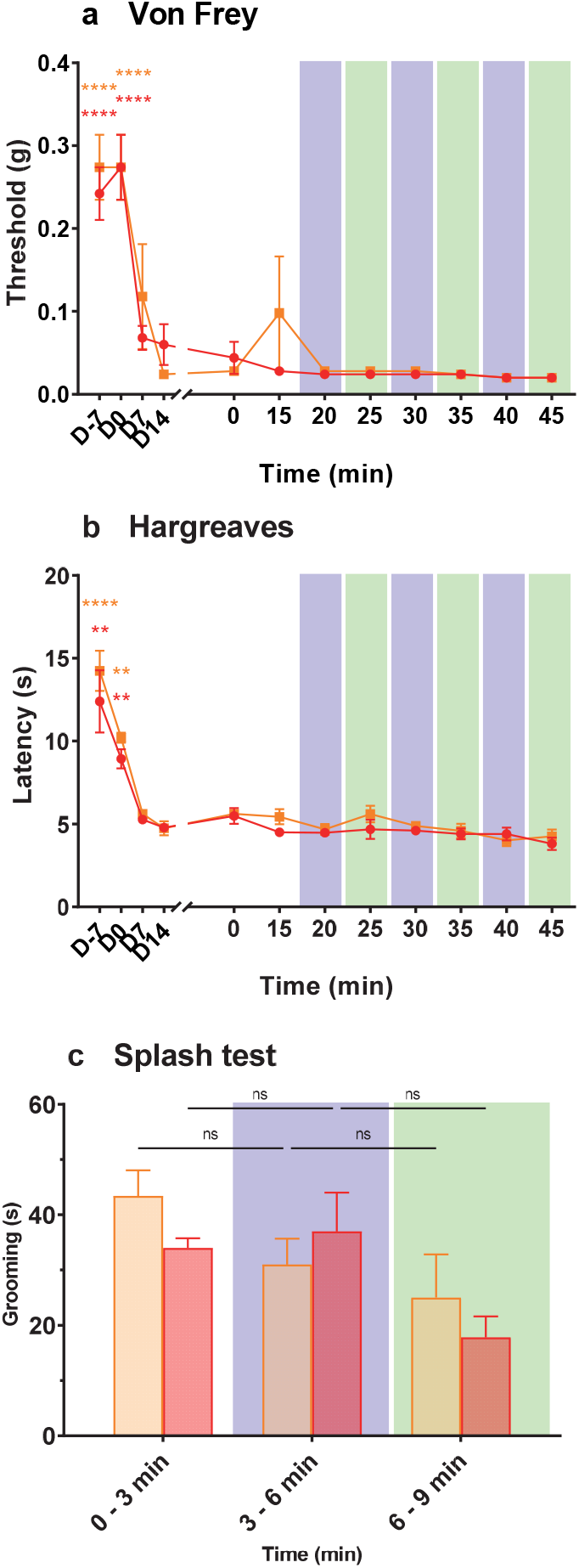
Lack of effect of UV/green light illumination in neuropathic mice injected by vehicle. Experiments were performed in the same conditions than those reported in Fig. 4, except that no photoswitchable ligand was injected. Green or violet light was delivered by a LED light source through an optic fiber connected to an hybrid optic-fluidic cannula implanted stereotaxically in the right or left amygdala of male mice with a cuff around the sciatic nerve of the left hind paw. **a-c:** Males, Cuff left hindpaw, vehicle (500 nL in PBS) delivered unilaterally in ipsi or controlateral amygdala. **a**. Von Frey. Mean ± SEM, t_x_ vs t_0_ (0 min), two-way ANOVA, Dunnett’s post-hoc test. **b**. Hargreaves. Mean ± SEM, t_x_ vs t_0_ (0 min), two-way ANOVA, Dunnett’s post-hoc test.. **c**. Splash Test. # t_x_ vs t_0_ (0 min), two-way ANOVA, Tukey’s post-hoc test.

**Supplemental figure 4:**
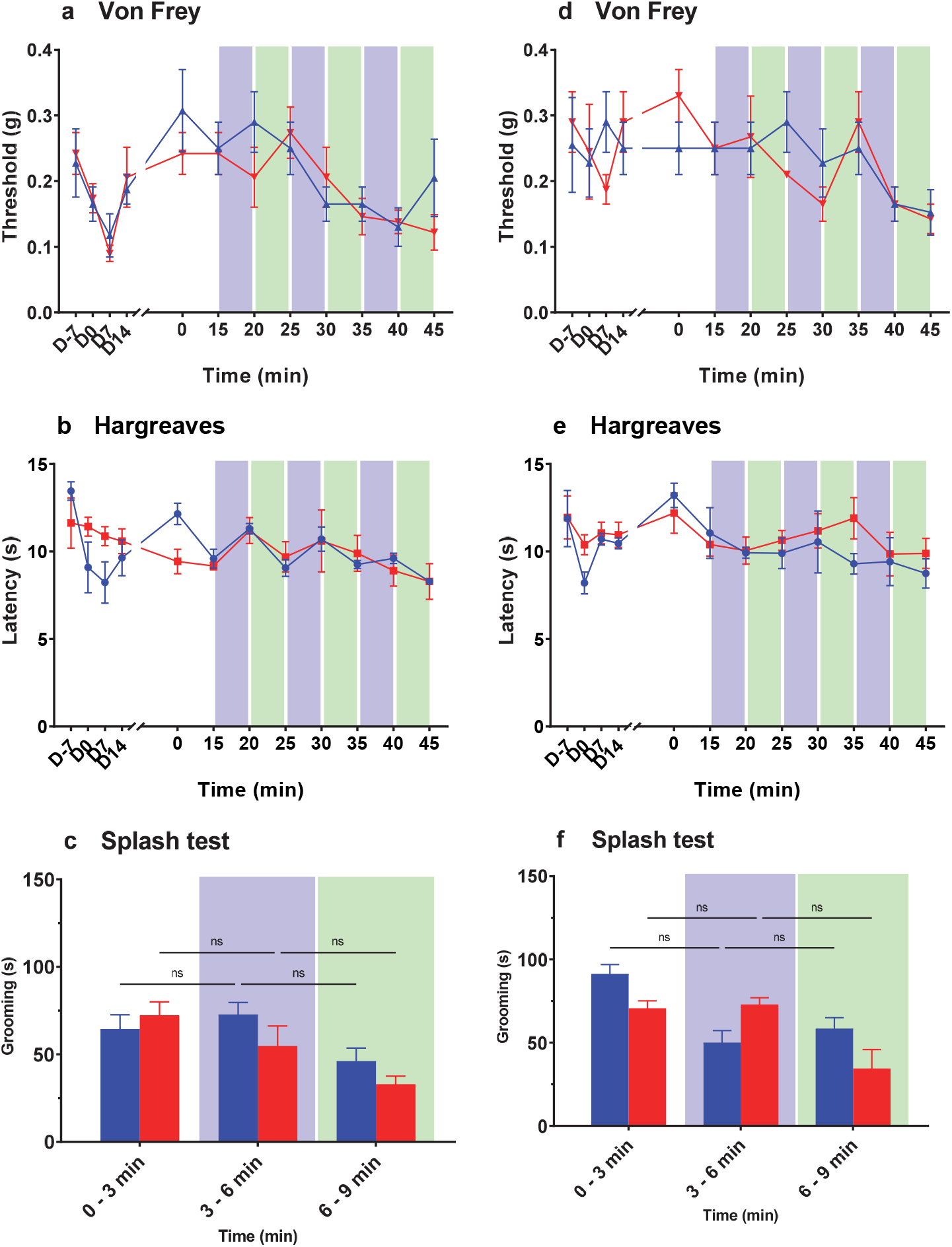
Photopharmacological manipulation of mGlu_4_ in right or left amygdala does not modify mechanical sensitivity, thermal sensitivity or grooming in Sham animals. **a-f:** Females, sham left hindpaw, Vehicle (500 nL PBS, blue) or LSP4-2022 (5 μM, 500 nL in PBS, red) delivered unilaterally in ipsi or controlateral amygdala. **a, d**. Von Frey. Mean ± SEM, t_x_ vs t_0_ (0 min), two-way ANOVA, Dunnett’s post-hoc test. **b, e**. Hargreaves. Mean ± SEM, t_x_ vs t_0_ (0 min), two-way ANOVA, Dunnett’s post-hoc test.. **c, f**. Splash Test. # t_x_ vs t_0_ (0 min), two-way ANOVA, Tukey’s post-hoc test.

**Supplemental figure 5:**
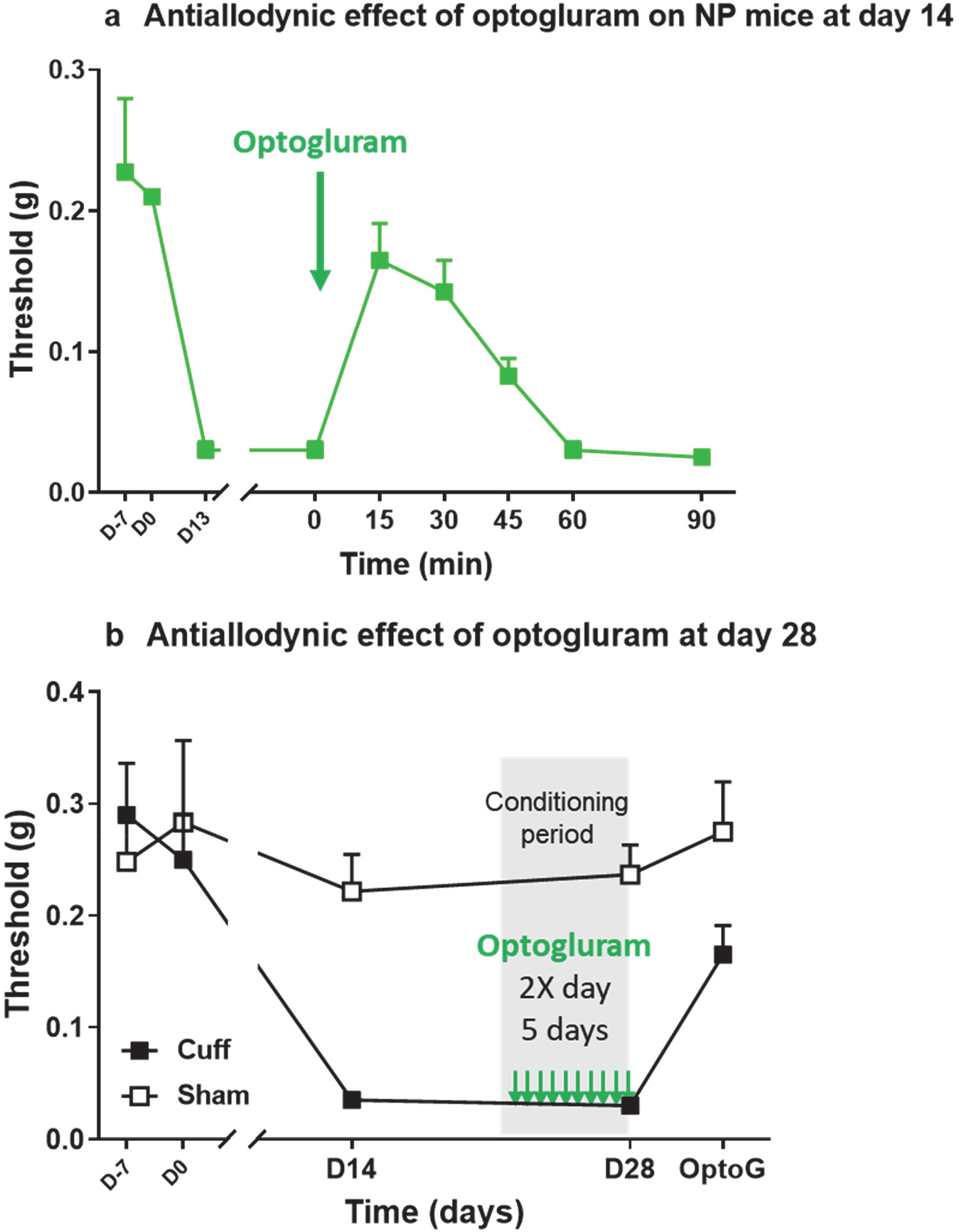
No tolerance following 5 days of chronic treatment (twice daily) of optogluram (30μM) during the conditioning period. Testing of aCPP 15 minutes following intra-amygdala injection of optogluram on day 6

**Supplemental figure 6:**
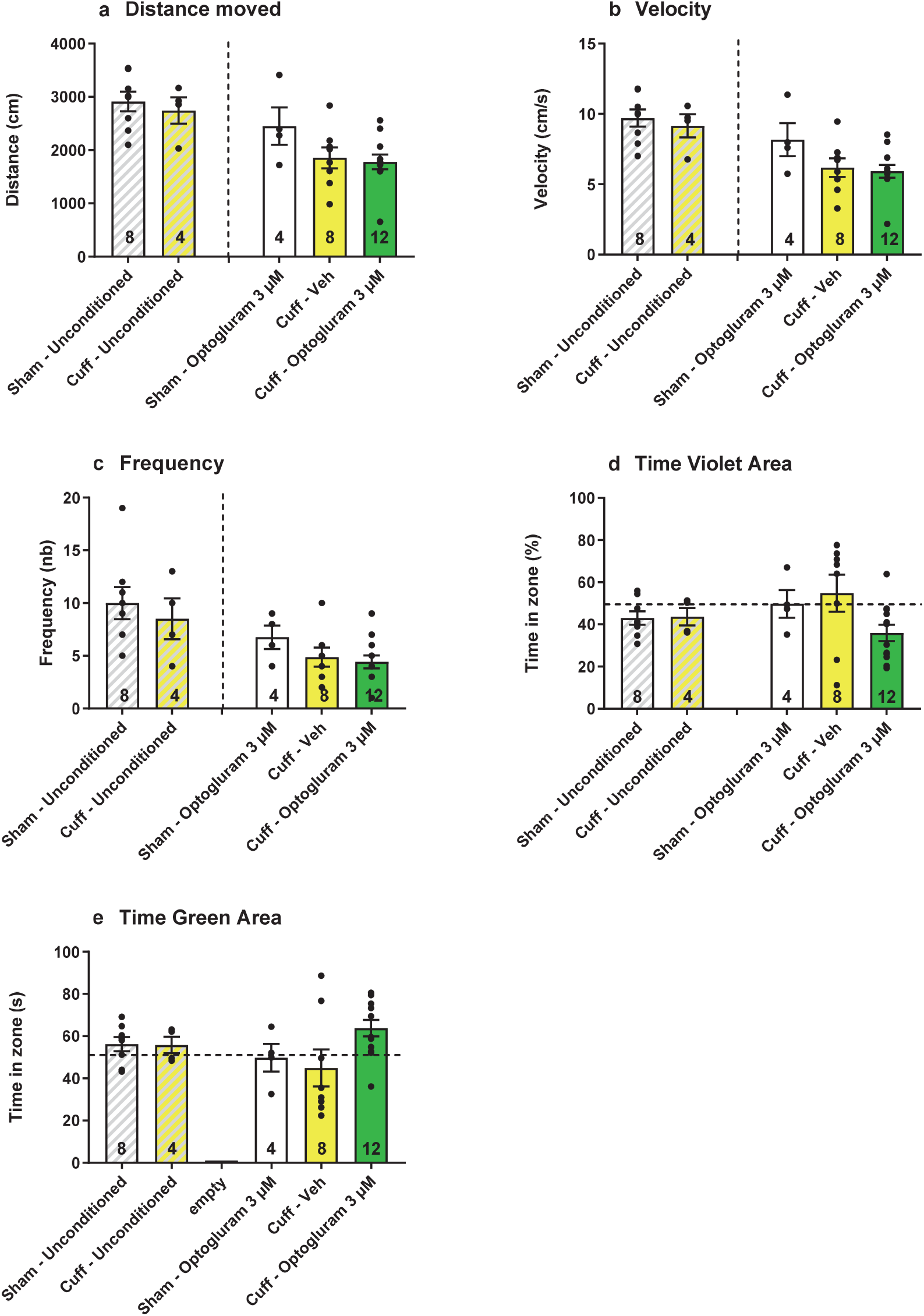
aCPP: different behavioural parameters on non-conditioned and conditioned Sham or neuropathic mice.

**Supplemental figure 7:**
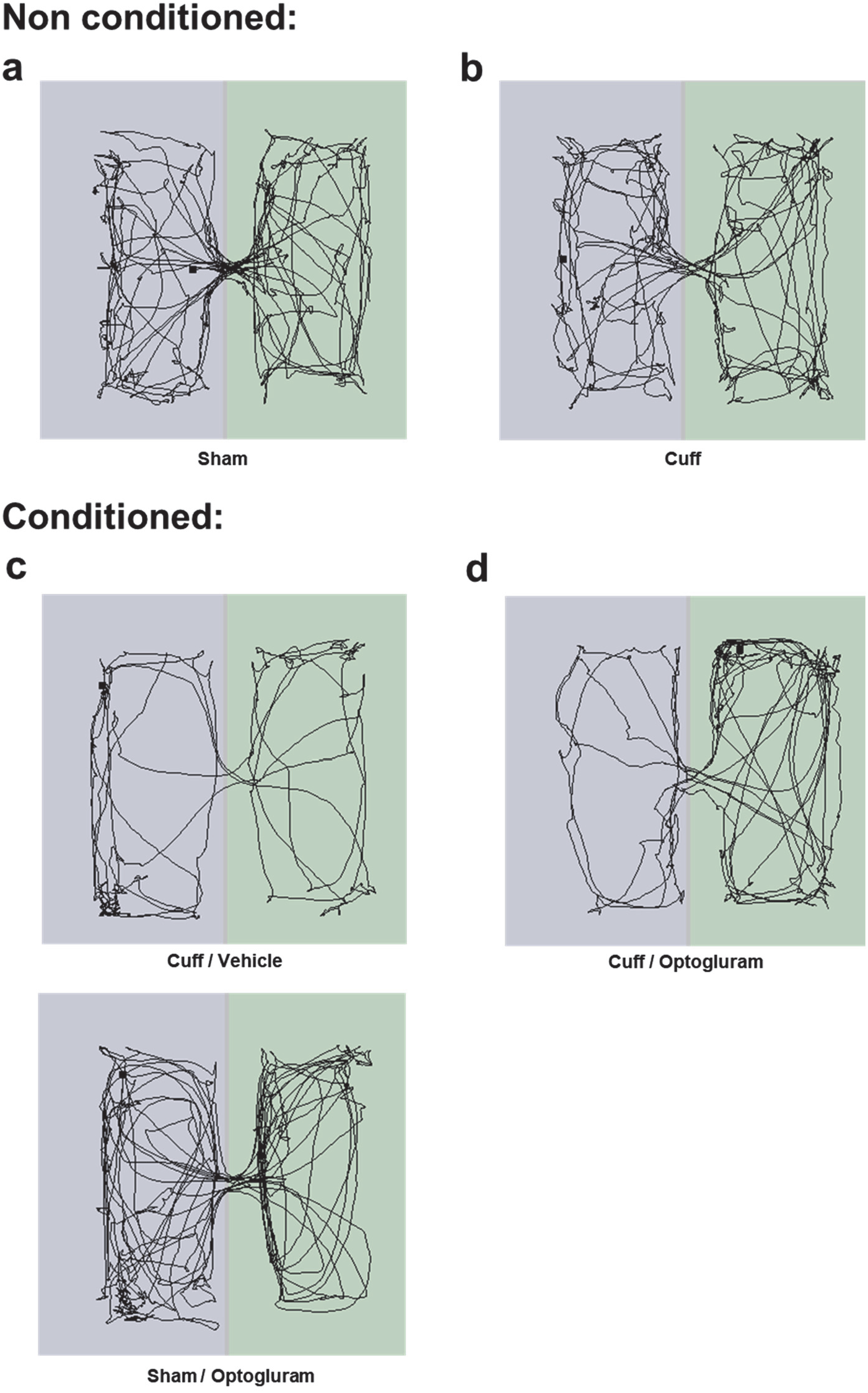
aCPP: Examples of videotracks on non-conditioned and conditioned Sham or neuropathic mice.

